# Squigulator: simulation of nanopore sequencing signal data with tunable noise parameters

**DOI:** 10.1101/2023.05.09.539953

**Authors:** Hasindu Gamaarachchi, James M. Ferguson, Hiruna Samarakoon, Kisaru Liyanage, Ira W. Deveson

## Abstract

*In silico* simulation of next-generation sequencing data is a technique used widely in the genomics field. However, there is currently a lack of optimal tools for creating simulated data from ‘third-generation’ nanopore sequencing devices, which measure DNA or RNA molecules in the form of time-series current signal data. Here, we introduce *Squigulator*, a fast and simple tool for simulation of realistic nanopore signal data. *Squigulator* takes a reference genome, transcriptome or read sequences and generates corresponding raw nanopore signal data. This is compatible with basecalling software from Oxford Nanopore Technologies (ONT) and other third-party tools, thereby providing a useful substrate for testing, debugging, validation and optimisation of nanopore analysis methods. The user may generate noise-free ‘ideal’ data, realistic data with noise profiles emulating specific ONT protocols, or they may deterministically modify noise parameters and other variables to shape the data to their needs. To highlight its utility, we use *Squigulator* to model the degree to which different types of noise impact the accuracy of ONT basecalling and downstream variant detection, revealing new insights into the properties of ONT data. We provide *Squigulator* as an open-source tool for the nanopore community: https://github.com/hasindu2008/squigulator

## INTRODUCTION

Nanopore sequencing is an increasingly important genomic technology. Devices from Oxford Nanopore Technologies (ONT) have the ability to analyse both short and long native DNA and RNA molecules, with countless potential applications across the life sciences. An ONT device measures the displacement of ionic current as a DNA or RNA molecule passes through a nanoscale protein pore. The device records time-series current signal data – commonly referred to as ‘squiggle’ data – which can be ‘basecalled’ into DNA/RNA sequence reads or analysed directly in a variety of contexts^1^.

Data simulation is an essential tool for data scientists and software developers in many scientific domains, including genomics^2^. The availability of reference data generated *in silico*, ideally with known error and/or noise profiles, enables the user to test, debug, optimise and validate new analysis methods in absence of confounding experimental and biological variables. Simulated data can also be used to develop and test new hypotheses or models, to inform experimental design, or as a ground truth during benchmarking studies, among other applications^2^. There are a range of existing tools for next-generation sequencing (NGS) data simulation, including *ART*^3^, *GemSIM*^4^, *pIRS*^5^ and *FASTQSim*^6^. These have helped catalyse important developments across genomics, transcriptomics, metagenomics, etc^2^. However, these popular tools generate sequence reads only and cannot be used to create realistic nanopore sequencing data with accompanying raw signal.

Here, we introduce *Squigulator* (<u>**Squig**</u>gle sim<u>**ulator**</u>), a fast and simple tool for *in silico* generation of nanopore current signal data that emulates the properties of real data from a nanopore device. The simulated data is compatible with ONT basecalling software and open-source tools for signal data analysis, providing a useful substrate for developers and data scientists in the growing nanopore community. *Squigulator* utilises existing ONT pore models and applies gaussian noise functions to generate realistic signal data from a reference sequence/s. The user can modulate these noise parameters, and other pseudo-experimental variables, such as the coverage depth and read-length distribution, to determine the properties of their simulated dataset. They may opt to create ‘ideal’ signal data lacking any noise, realistic noisy data, or instead model the impact of one or more noise parameters on the performance of their analysis workflow. This capacity for deterministic control of noise parameters is an important advantage of *Squigulator*, enabling parameter exploration during algorithm development.

## METHODS & IMPLEMENTATION

*Squigulator* generates simulated nanopore signal data based on an input reference genome or transcriptome sequence (FASTA format), or directly from a set of basecalled reads (FASTQ or FASTA format). To do so, it uses an idealised ‘pore model’ that specifies the predicted current signal reading associated with every possible DNA or RNA *k*-mer, as appropriate to the specific nanopore protocol being emulated. Pore models are available for ONT protocols (https://github.com/nanoporetech/kmer_models), covering a range of different flow-cells (e.g. R9, R10), kits (e.g. DNA vs RNA sequencing) and ONT devices (e.g. PromethION vs MinION). The user can also provide their own custom pore model, enabling simulation of data from existing or future non-ONT nanopore systems.

*Squigulator* generates sequential signal values corresponding to sequential *k*-mers in the provided reference sequence. The number of signal values per *k*-mer is dictated by the translocation speed for a specified ONT sequencing protocol (e.g. 450 nt/sec. for DNA sequencing on R9.4.1 flow cells) or provided manually by the user. This process generates perfect, noise-free signal reads when the *--ideal* option is selected (**Fig1**). By default, however, *Squigulator* transforms the data using Gaussian noise functions in both the time and amplitude domains to produce realistic, rather than ideal, signal reads (**Fig1**). The user can select from a menu of pre-set configurations with noise parameters and other variables empirically modelled to emulate specific ONT protocols. Alternatively, they can use the *--ideal-time* or *--ideal-amp* options to limit noise to just a single domain, or they can manually adjust both noise parameters independently (**Fig1**). The user can also modify several pseudo-experimental variables, including translocation speed, coverage depth, read-length mean, variation, etc, to determine the dimensions of their simulated dataset.

**Fig1.**
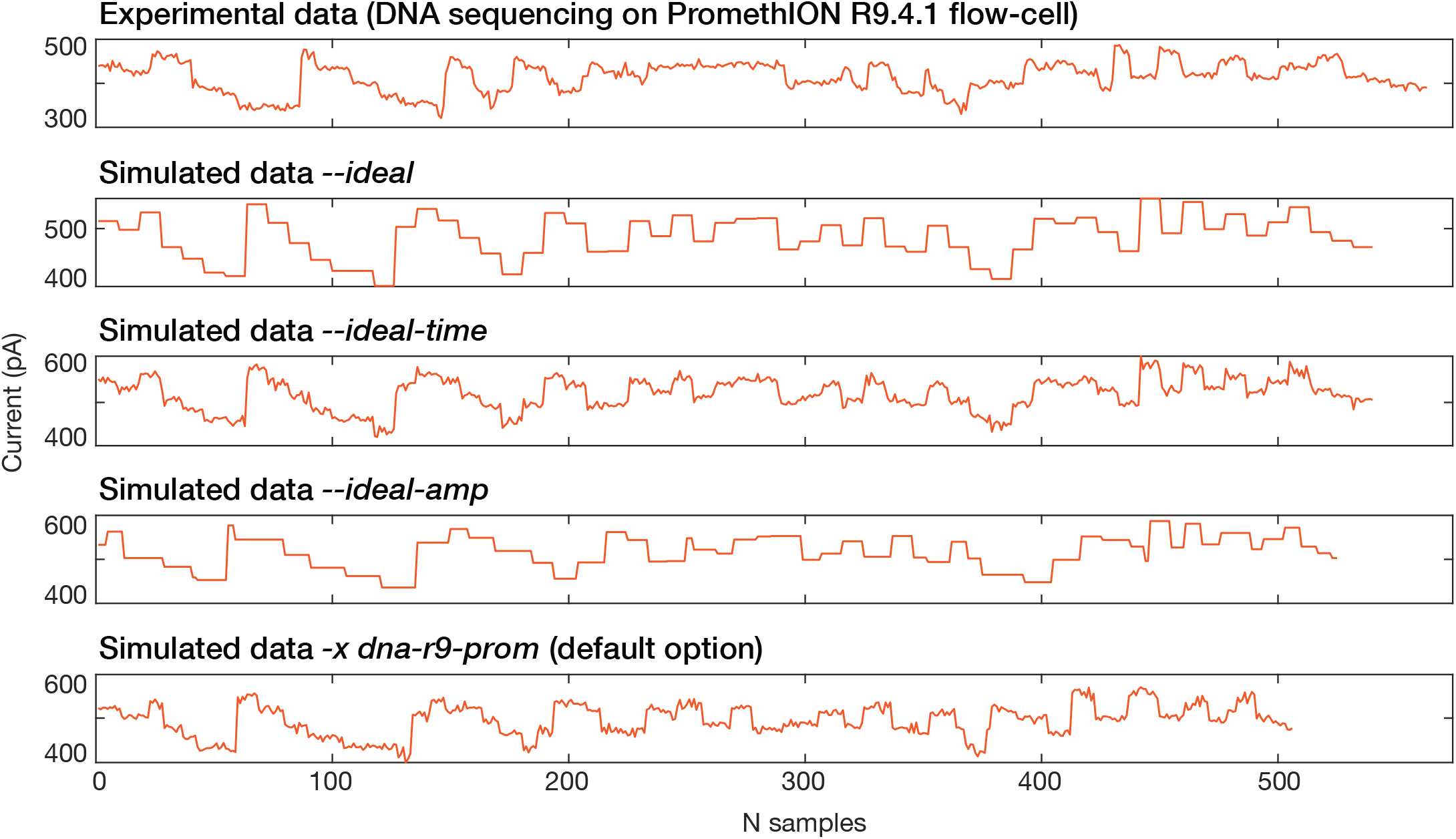
Simulation nanopore signal data with tunable noise parameters. This example compares experimental nanopore raw signal data (top track) to corresponding simulated data from *Squigulator* with various noise parameter settings. The experimental data was generated by sequencing human genomic DNA on an ONT PromethION R9.4.1 flow cell. The realistic simulated data (bottom track) emulates this by using the *-x dna-r9-prom* pre-set configuration, which is currently the default option in *Squigulator*.

*Squigulator* is developed in C programming language. The only external dependency of *Squigulator* is the ubiquitous library *zlib*. The *Slow5lib* library^7^ for reading and writing nanopore signal data is included inside the source code of *Squigulator* itself. The minimum compiler requirement is a C99 or higher C compiler that supports POSIX extensions (e.g., *gcc, clang* or *icc*). Therefore, the software can be easily compiled on Linux, MacOS and Windows (through Windows Subsystem for Linux).

The core random number generator used by *Squigulator* is a simple uniform random number generator based on the simple Multiplicative Linear Congruential Generator (LCG). This provides the basis for generating normal distributions (using Box-Muller transform) and gamma distributions, where appropriate. Sampling the genome at random positions for reads is modelled as a uniform random distribution. Normal distributions are used for modelling noise along amplitude domains of the signal, where the mean and standard deviation of each *k*-mer in the pore-model form a random number generator. Normal distributions are also used for generating noise along the time domain (dwell), and other signal metadata such as *offset* and *median_before*. Read length variability is modelled using a gamma distribution. In each case, the most appropriate distribution type was selected based on empirical exploration of real experimental ONT data.

## RESULTS

### Squigulator compatibility with nanopore analysis software

*Squigulator* is designed to produce simulated nanopore signal data that is compatible with any relevant software and can be used to recapitulate a complete nanopore analysis workflow. To test this capability, we created a dataset that resembles a typical sequencing experiment run on the popular human reference individual NA12878 (see **Supplementary Methods**). Briefly, we used *bcftools consensus*^8^ to incorporate high-confidence NA12878 variants (SNVs and indels) annotated by the Genome in a Bottle consortium^9^ into the human reference genome sequence (*hg38*; FASTA format). This was done in a haplotype-aware fashion, creating a diploid NA12878 reference in which heterozygous and homozygous variants are represented at appropriate copy number. We then used *Squigulator* to generate simulated nanopore signal data from this custom reference, with default noise parameters. The read-length, standard deviation and read-count were roughly matched to a real sequencing experiment performed on NA12878 genomic DNA (∼30X coverage; see **Supplementary Methods**), which is used for comparison below.

We analysed both the simulated and experimental NA12878 datasets via a typical analysis workflow. Signal data was basecalled with ONT’s *Guppy* software (using the *Buttery-eel* wrapper for SLOW5 data access)^10^. Basecalled reads were then aligned to the reference genome using *Minimap2*^11^ and single-nucleotide variants (SNVs) were detected using each of two approaches: (i) with *Nanopolish*^12^, which uses both the basecalled and raw-signal data to identify variants and; (ii) with *Clair3*^13^, which calls variants from the basecalled data alone. SNV detection performance was then evaluated with *rtgtools vcfeval* (see **Supplementary Methods**).

We observed broadly similar properties between the simulated and experimental NA12878 datasets at each workflow stage. The distribution of signal values were similar between the two raw datasets (**Fig2a**). After basecalling, they showed similar read-length, nucleotide composition and quality score distributions (**Fig2b,c**). *Minimap2* alignment statistics were similar (**Supplementary Table1**), as were the patterns of mismatch and indel errors in reads aligned to hg38 (**Fig2d,e**). Both variant detection tools were able to detect known NA12878 SNVs with accuracy over 98% (F-score), with *Clair3* showing superior performance to *Nanopolish* on both the simulated and experimental datasets (**Supplementary Table2**). These results demonstrate the capacity of *Squigulator* to create simulated data that emulates real experimental data and is compatible with relevant analysis software.

**Fig2.**
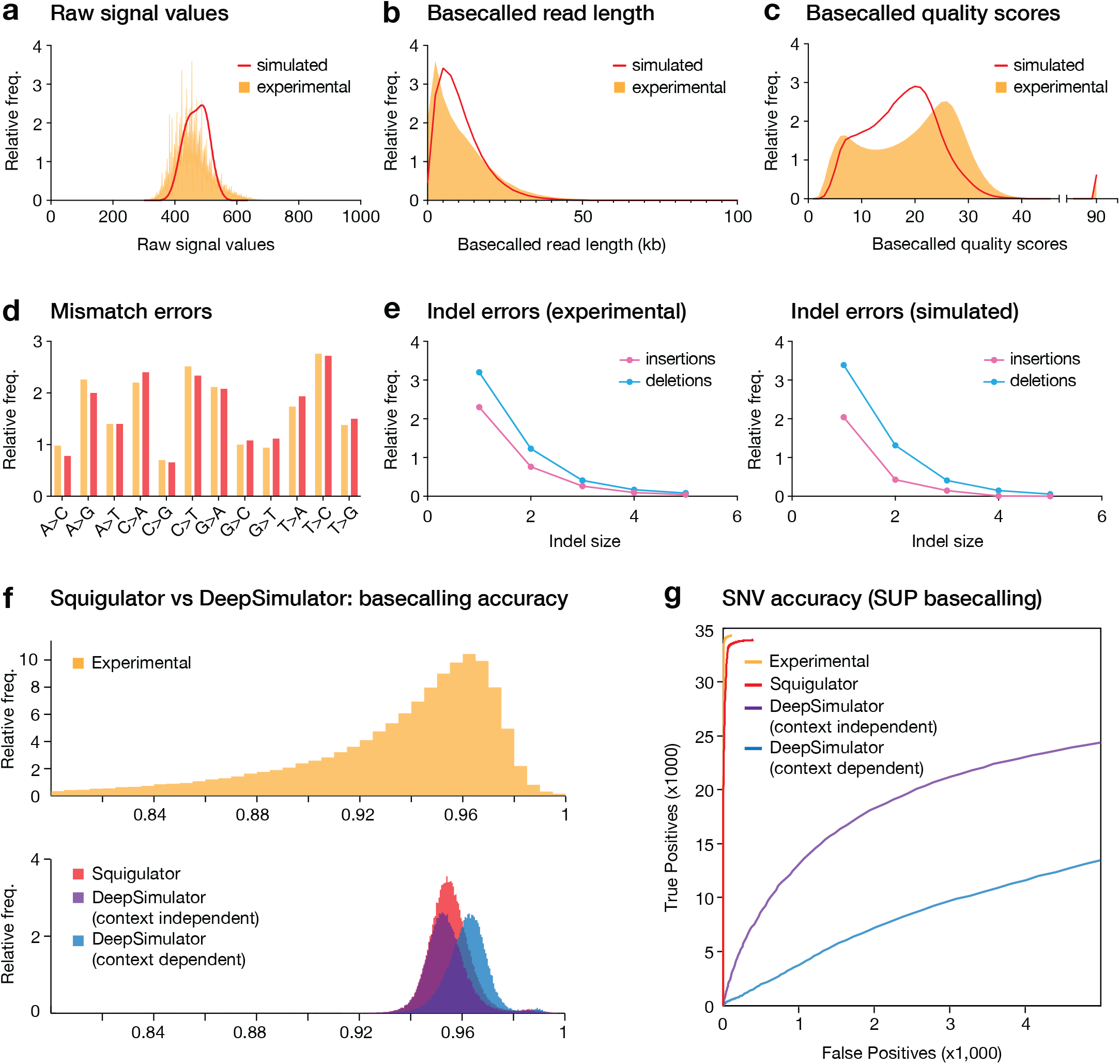
Comparison of experimental and simulated NA12878 signal datasets. (**a-c**) Frequency histograms show distributions of (**a**) raw signal values, (**b**) basecalled read lengths, and (**c**) Phred quality scores in experimental (orange) and simulated (red) datasets, based on the reference individual NA12878. Guppy HAC basecalling model was used. (**d**,**e**) For the same datasets, bar chart show the relative frequencies of each possible base substitution (**d**) and line plots show the relative frequency of insertions (pink) and deletions (blue) of different sizes (**e**). Substitution and indels errors are determined relative to the hg38 reference genome after alignment with *Minimap2*. (**f**) *Guppy* basecalling accuracy (HAC model), as measured by read:reference identity score distributions, for experimental (upper) and simulated (lower) datasets. Simulated data is from *Squigulator* (red) or *DeepSimulator* with context-independent (purple) or context-dependent (blue) settings. (**g**) ROC curves evaluate accuracy of SNV detection by *Clair3* on the same datasets (colours are matched). SUP basecalling was used to maximise accuracy of SNV detection.

### Comparison of Squigulator to DeepSimulator

There are currently two mature tools available for nanopore data simulation: *NanoSim*^14^ and *DeepSimulator*^15^. *NanoSim* generates basecalled sequencing data (FASTQ format) but does not generate current signal data, and is therefore not suitable for comparison to *Squigulator. DeepSimulator* generates simulated signal data via either of two approaches: (i) using a context-dependent Bi-LSTM trained model or; (ii) a context-independent model that utilises a *k*-mer model provided by ONT. Both approaches are designed to generate realistic simulated data that faithfully emulates real nanopore signal data, though the context-independent model is considerably faster.

To compare the performance of *Squigulator* and *DeepSimulator*, we repeated the experiment above but this time generating simulated NA12878 datasets using *DeepSimlulator’s* context-dependent and context-independent modes (see **Supplementary Methods**). We assessed the accuracy of *Guppy* base-calling and *Clair3* SNV detection for each simulated dataset, with the real experimental NA12878 dataset as a point of comparison. While the differences were relatively modest, *Squigulator* basecalled data showed higher accuracy (median 0.948 read:reference identity) than *DeepSimulator*’s context-independent mode (0.946) and lower accuracy than the context-dependent mode (0.958; *Guppy* HAC model; **Fig2f**). All simulated datasets showed broadly similar accuracy to real experimental data. However, inspecting alignments to the reference genome, we observed that basecalled data from *DeepSimulator* exhibited many systematic errors, whereas errors in *Squigulator* data and experimental data were distributed more randomly (**FigS1**). As a result, *Clair3* called an abundance of erroneous SNVs on both *DeepSimulator* datasets and missed many true SNVs. Overall NA12878 SNV accuracy with *DeepSimulator* data (context-dependent 0.625; context-independent 0.769 F-scores; *Guppy* SUP) was poor by comparison to *Squigulator* (0.987) or real experimental data (0.996; **Fig2g**).

In addition to data quality, we compared the time and resources used by the two simulation tools. Running with 16 CPUs, *Squigulator* took just 156 seconds to generate ∼30X sequencing data on chr22 (with peak RAM usage of 0.5 GB) or 68 minutes for an entire human genome (3.4GB RAM; **Supplementary Table 3**). To generate an equivalent chr22 dataset on the same system, *DeepSimulator* took ∼20 times longer (50 minutes) in context-independent mode and ∼3000 times longer (131 hours) in context-dependent mode (**Supplementary Table 3**). *DeepSimulator’s* peak RAM usage was also ∼9-fold and ∼206-fold higher than *Squigulator*, respectively. While these analyses show that *DeepSimulator* is able to generate simulated nanopore signal data, *Squigulator* requires a fraction of the time, memory, and generates data that more closely resembles real experimental data.

### Using Squigulator for parameter exploration

While they showed similar patterns, we observed higher overall error rates during basecalling and SNV detection with the simulated vs experimental datasets above (**Supplementary Table 1**). Anticipating that these processes may be affected by noise in the time and amplitude domains, we next used *Squigulator* to model the impact of each noise parameter. To do so we generated alternative simulated datasets in which the degree of noise in one domain was systematically modified, while keeping the other static, then repeated the above analysis workflow with each dataset (see **Supplementary Methods**).

Both noise parameters had a significant impact on basecalling accuracy. Accuracy was negatively correlated with noise in the time domain (**Fig3a**) and surpassed real experimental data when noise was minimised (**FigS2a**). In fact, basecalling was much more sensitive to dwell-time noise (i.e. standard deviation) than to fixed changes in the dwell-time mean (**FigS2b,c**). This indicates that consistency in DNA translocation speed is more important to *Guppy* than the actual speed itself (within a sensible range). Noise in the amplitude domain was also detrimental to basecalling accuracy, however, reducing amplitude noise to zero did not lead to optimum results (**Fig3b**). Instead, *Guppy* performed best when a small amount of noise applied, presumably reflecting properties of the neural network models involved, which are trained on real experimental data. Interestingly, these trends were not consistent between *Guppy’s* FAST, HAC or SUP models, with HAC being more sensitive to changes in the degree of amplitude noise than SUP or FAST (**Fig3b**). As expected, differences in basecalling accuracy at the read-level resulting from noise in the time and amplitude domains manifested in corresponding differences in the accuracy of SNV detection *Clair3* (**Fig3b,c**). Overall, this analysis provides a simple demonstration of how simulated data with tunable noise parameters may be used to inform algorithm development and optimisation in the nanopore field.

**Fig3.**
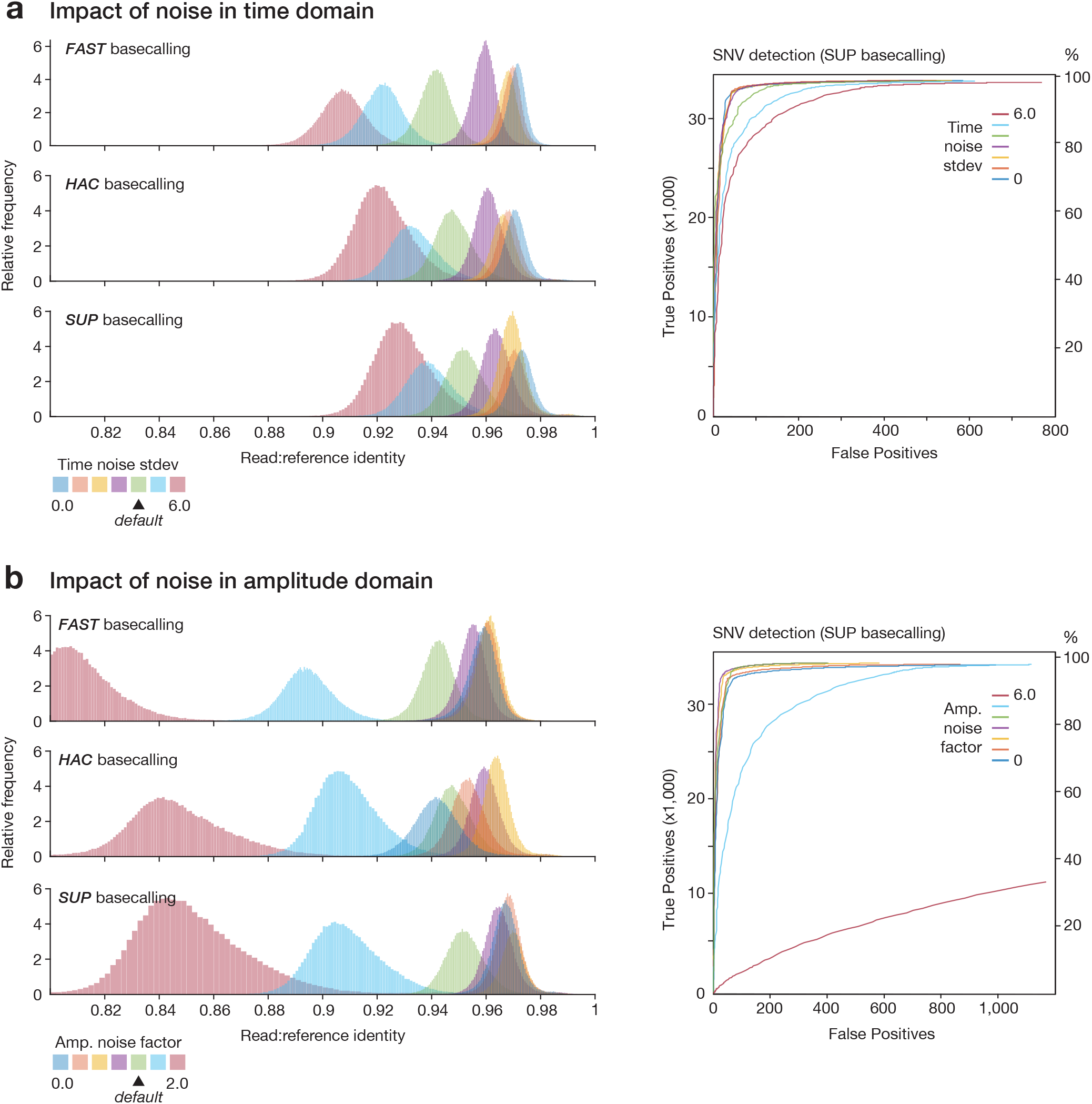
Modelling the impact of time and amplitude noise on sequencing accuracy. (**a**; left) *Guppy* basecalling accuracy, as measured by read:reference identity score distributions, for repeated experiments in which noise in the time domain (--dwell-std) is varied, while other parameters are held at default. Higher values indicate increasing noise and default value is --dwell-std=4 (green distribution). Experiment was repeated with *Guppy’s* FAST (upper), HAC (middle) and SUP (lower) basecalling models. (**a**; right) ROC curves assess accuracy of SNV detection by *Clair3* on the same datasets (colours are matched). (**b**) Same analysis as above but with noise in the amplitude domain (--amp-noise) varied while other parameters are static.

## DISCUSSION

The genomics field has benefitted from the availability of many different tools for *in silico* data simulation, which have been developed in parallel to new sequencing technologies and applications^2^. *Squigulator* addresses the need for a simple and effective tool for creating realistic simulated data from nanopore sequencing experiments.

The most popular tool currently available for nanopore data simulation, *NanoSim*^14^ generates basecalled sequencing data (FASTQ format) but does not generate current signal data, which is the primary output from a nanopore device. Therefore, *NanoSim* cannot be used to recapitulate a complete nanopore analysis workflow, and has no utility for development and benchmarking of signal-level analyses, such as basecalling. In contrast, we have shown above how *Squigulator* can be used to generate realistic signal data, suitable to evaluate basecalling and all relevant downstream tools.

Another alternative, *DeepSimulator*^15^, uses a neural network architecture to generate realistic nanopore signal data. This approach is computationally expensive but is designed to best emulate the subtleties of experimental data on which it was modelled. In contrast, *Squigulator’s* simple, deterministic framework is designed to be fast, lightweight and grant the user maximum control over all variables and noise parameters. Surprisingly, our comparison experiments show that simulated data from *Squigulator* is more similar to matched experimental data than can be achieved with either *DeepSimulator’s* context-dependent or context-independent modes. Although *Guppy* basecalling accuracy was relatively similar overall, *DeepSimulator* data exhibited systematic errors that produced an abundance of false-positives during SNV detection with *Clair3*.

Generating realistic data is not the limit of *Squigulator’s* capabilities. The user can instead tune experimental variables and noise parameters to assess the impact on their analysis workflow. The pore-model input framework allows the user to select from pre-existing pore models and pre-set noise parameters for different ONT protocols, or can provide their own custom pore model and/or parameters. This means *Squigulator* is, in theory, suitable for simulating data from any alternative nanopore system where the user has access to a *k*-mer-based pore model (e.g. a hypothetical solid state nanopore device with voltage sensing). Therefore, while both simulation tools can generate useful nanopore signal data, *Squigulator* provides key advantages in data quality and parameter control over *DeepSimulator*.

An effective tool for *in silico* data generation should reduce friction in bioinformatics software development. Speed and simplicity are essential to achieve this aim. Therefore, *Squigulator* is designed to be easy to install and use. It has no external dependencies apart from the ubiquitous *zlib* software library and can be compiled on Linux, MacOS and Windows, or simply run using pre-compiled binaries provided for common systems. *Squigulator* is also exceptionally fast by comparison to other similar tools. We have found that generating 1Gbase of simulated data takes around 2 minutes when running on 16 CPU threads, compared to ∼45 minutes with *DeepSimulator’s* low-accuracy context-independent mode and ∼90 hours with its context-dependent mode (a comparison with *NanoSim* was not possible because we were unable to complete the installation). This combination of speed, simplicity and realistic, tunable data (see above) make *Squigulator* the ideal tool for nanopore data simulation.

Squigulator outputs nanopore signal data in binary SLOW5 (BLOW5) format, an open source community-centric alternative to ONT’s native FAST5 format^7^. This feature is essential to the tool’s high performance, since the writing speed for BLOW5 is considerably faster than what is attainable with FAST5. BLOW5 format is now compatible with the latest ONT basecalling models and approaches, via *Buttery-eel* (a SLOW5 wrapper for ONT’s production basecaller, *Guppy*)^10^ or SLOW5-enabled forks for open source basecallers *Dorado* (https://github.com/hiruna72/slow5-dorado) or *Bonito* (https://github.com/Psy-Fer/bonito). BLOW5 format is also compatible with a wide variety of open source software^16–21^. Users with masochistic tendencies may alternatively decide to convert their BLOW5 files back to ONT’s native FAST5 format using *slow5tools*^*22*^, before proceeding to their analysis workflow.

## DATA & CODE AVAILABILITY

The experimental dataset used in benchmarking experiments is available on NCBI Sequence Read Archive at Bioproject PRJNA744329. All other benchmarking datasets can be created using *Squigulator* by following instructions in the **Supplementary Methods** section. With the exception of ONT’s commercially available *Guppy* basecaller, all software used in this project is free and open source, including *Squigulator*: https://github.com/hasindu2008/squigulator Squigulator experiments were performed using the following github commit: 7422d7384be428ac334caa61c019473f31f1e633

## ACKNOWLEDGEMENTS

We thank Derrick Lin and Tim Ho for providing HPC support. We acknowledge the following funding support: Australian Medical Research Futures Fund grants MRF1173594, MRF2016008 and MRF2023126 (to I.W.D.) and Australian Research Council DECRA Fellowship DE230100178 (to H.G.).

## DECLARATIONS

I.W.D. manages a fee-for-service sequencing facility at the Garvan Institute of Medical Research that is a customer of Oxford Nanopore Technologies but has no further financial relationship. H.G., J.M.F. and I.W.D. have previously received travel and accommodation expenses to speak at Oxford Nanopore Technologies conferences. H.G. and I.W.D. have paid consultant roles with Sequin PTY LTD. The authors declare no other competing financial or non-financial interests.

## CONTRIBUTIONS

H.G. developed *Squigulator* with contributions from all other authors. H.S., K.L., and J.M.F conducted rigorous user testing and feedback. I.W.D., H.G. and J.M.F. performed benchmarking experiments. H.G. and I.W.D. generated the figures and wrote the manuscript, with input from other authors.

## FIGURE LEGENDS

**FigS1.**
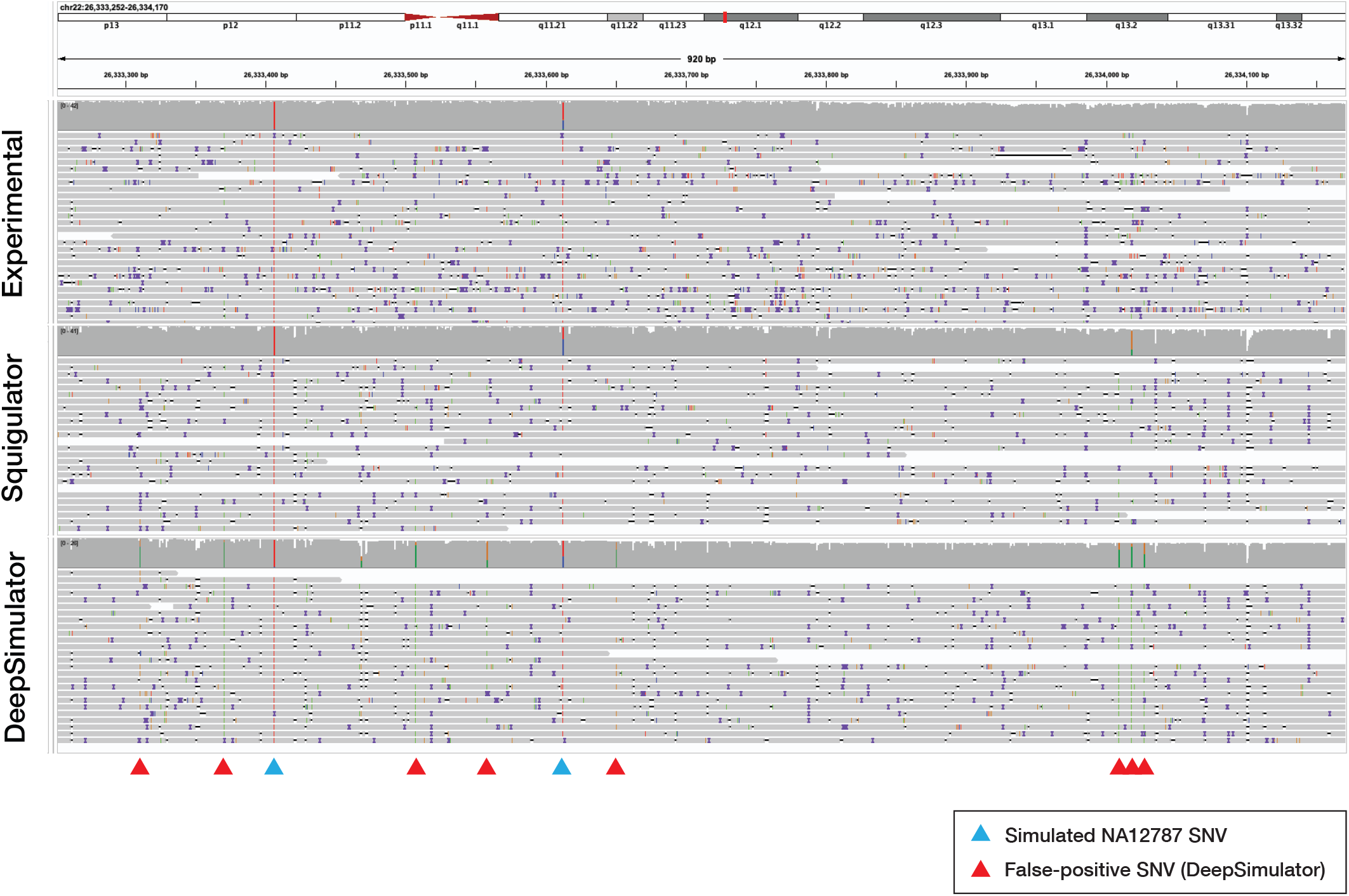
Comparison of Squigulator and DeepSimulator to real experimental ONT data. Genome browser view shows basecalled reads (Guppy SUP model) aligned to the human reference genome (hg38). The top track shows real experimental data from ONT sequencing of NA12878 genomic DNA (R9.4.1 PromethION flow cells). The middle track shows simulated NA12878 data from Squigulator with -x dna-r9-prom pre-set configuration. The bottom track shows simulated NA12878 data from DeepSimulator running in context-independent mode. Blue triangle markers show the location of NA12878 SNVs that were incorporated into the simulation, and are correctly detected by Clair3. Red triangle markers show the presence of systematic errors in basecalled reads from DeepSimulator, which are erroneously detected as SNVs by Clair3.

**FigS2.**
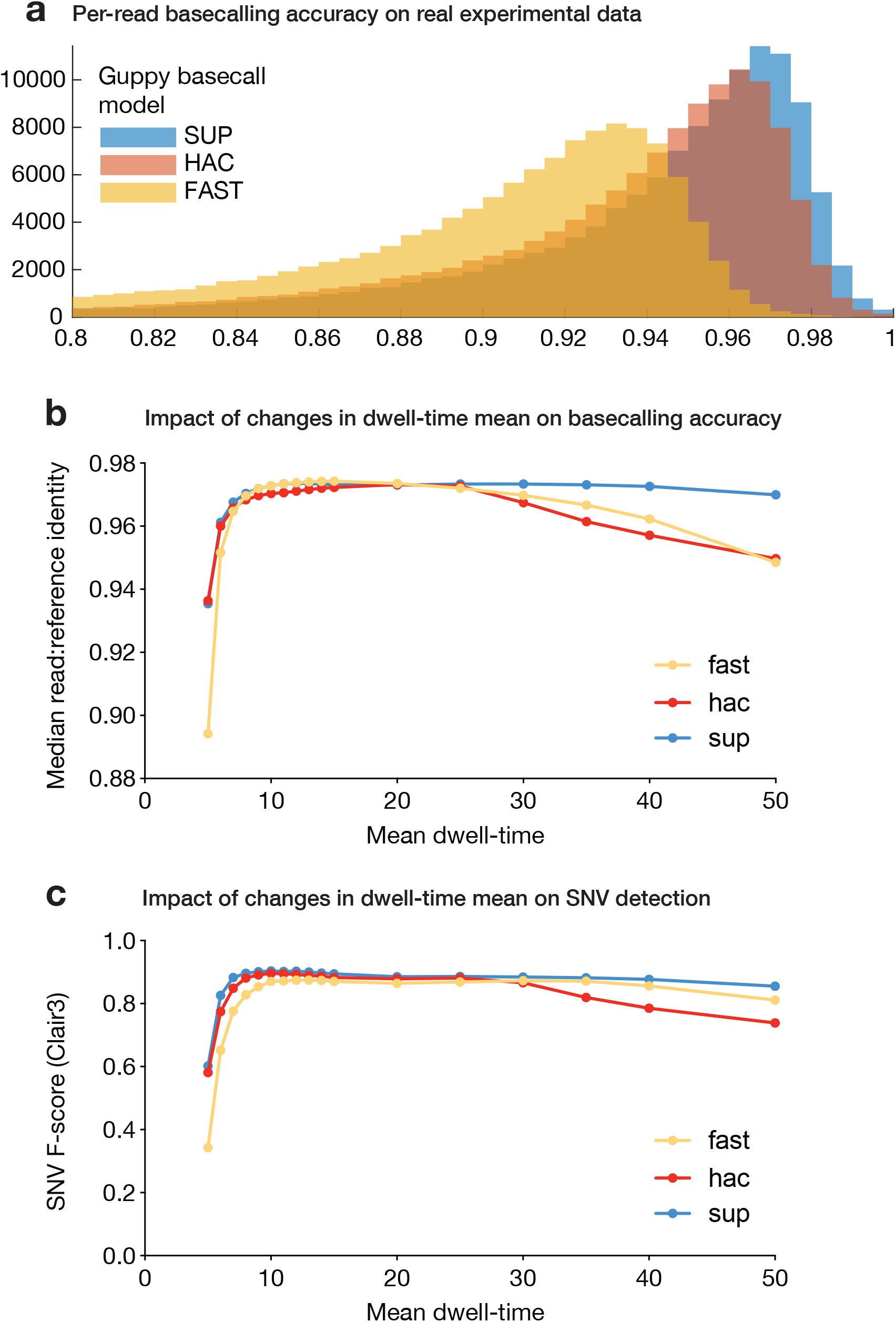
Parameter exploration regarding Guppy basecalling sequencing accuracy. (**a**) Guppy basecalling accuracy, as measured by read:reference identity score distributions, on real experimental NA12878 data with Guppy’s FAST, HAC or SUP models. (**b**) Guppy basecalling accuracy, as measured by read:reference identity score medians, for repeated experiments in which the mean dwell time (--dwell-mean) is varied, while other parameters are held at default. Experiment was repeated with FAST, HAC and SUP basecalling models. Default value --dwell-mean=9 (for R9.4.1 flow cell). (**c**) Accuracy of SNV detection, as measured by F-score, by Clair3 on the same datasets and basecalling models as above (colours are matched).

## REFERENCES

1. Wang, Y., Zhao, Y., Bollas, A., Wang, Y. & Au, K. F. Nanopore sequencing technology, bioinformatics and applications. Nat. Biotechnol. 39, 1348–1365 (2021).

2. Escalona, M., Rocha, S. & Posada, D. A comparison of tools for the simulation of genomic next-generation sequencing data. Nat. Rev. Genet. 17, 459–469 (2016).

3. Huang, W., Li, L., Myers, J. R. & Marth, G. T. ART: a next-generation sequencing read simulator. Bioinformatics 28, 593–594 (2012).

4. McElroy, K. E., Luciani, F. & Thomas, T. GemSIM: general, error-model based simulator of next-generation sequencing data. BMC Genomics 13, 74 (2012).

5. Hu, X. et al. pIRS: Profile-based Illumina pair-end reads simulator. Bioinformatics 28, 1533–1535 (2012).

6. Shcherbina, A. FASTQSim: platform-independent data characterization and in silico read generation for NGS datasets. BMC Res. Notes 7, 533 (2014).

7. Gamaarachchi, H. et al. Fast nanopore sequencing data analysis with SLOW5. Nat. Biotechnol. 40, 1026–1029 (2022).

8. Danecek, P. et al. Twelve years of SAMtools and BCFtools. Gigascience 10, (2021).

9. Zook, J. M. et al. Integrating human sequence data sets provides a resource of benchmark SNP and indel genotype calls. Nat. Biotechnol. 32, 246–251 (2014).

10. Samarakoon, H., Ferguson, J. M., Gamaarachchi, H. & Deveson, I. W. Accelerated nanopore basecalling with SLOW5 data format. bioRxiv (2023) doi:10.1101/2023.02.06.527365. [in press]

11. Li, H. Minimap2: pairwise alignment for nucleotide sequences. Bioinformatics 34, 3094–3100 (2018).

12. Loman, N. J., Quick, J. & Simpson, J. T. A complete bacterial genome assembled de novo using only nanopore sequencing data. Nat. Methods 12, 733–735 (2015).

13. Zheng, Z. et al. Symphonizing pileup and full-alignment for deep learning-based long-read variant calling. Nat. Comp. Sci. 2, 797–803 (2022).

14. Yang, C., Chu, J., Warren, R. L. & Birol, I. NanoSim: nanopore sequence read simulator based on statistical characterization. Gigascience 6, 1–6 (2017).

15. Li, Y. et al. DeepSimulator1.5: a more powerful, quicker and lighter simulator for Nanopore sequencing. Bioinformatics 36, 2578–2580 (2020).

16. Senanayake, A., Gamaarachchi, H., Herath, D. & Ragel, R. DeepSelectNet: deep neural network based selective sequencing for oxford nanopore sequencing. BMC Bioinformatics 24, 31 (2023).

17. Simpson, J. T. et al. Detecting DNA cytosine methylation using nanopore sequencing. Nat. Methods 14, 407–410 (2017).

18. Zhang, H. et al. Real-time mapping of nanopore raw signals. Bioinformatics 37, i477–i483 (2021).

19. Bao, Y. et al. SquiggleNet: real-time, direct classification of nanopore signals. Genome Biol. 22, 298 (2021).

20. Gamaarachchi, H. et al. GPU accelerated adaptive banded event alignment for rapid comparative nanopore signal analysis. BMC Bioinformatics 21, 343 (2020).

21. Shih, P.J. et al. Efficient Real-Time Selective Genome Sequencing on Resource-Constrained Devices. arXiv (022) doi:10.48550/arXiv.2211.07340.

22. Samarakoon, H. et al. Flexible and efficient handling of nanopore sequencing signal data with slow5tools. Genome Biol. 24, 69 (2023).

