## Supplementary Information for "Squigulator: simulation of nanopore sequencing signal data with tunable noise parameters"

### SUPPLEMENTARY METHODS

***Generating the experimental NA12878 dataset***

The experimental dataset used in benchmarking experiments was generated by sequencing genomic DNA from the human NA12878 reference sample on an ONT PromethION device. Unsheared DNA libraries were prepared using the ONT LSK109 ligation library prep, and two R9.4.1 flow-cells were used to generate ~30× genome coverage. The data is available on NCBI Sequence Read Archive at Bioproject PRJNA744329.

***Generating the simulated NA12878 dataset***

The simulated NA1278 dataset was generated using *Squigulator*, with the intention to emulate the real experimental dataset above. *Bcftools consensus* (v1.16) was used to incorporate high-confidence NA12878 variants (SNVs and indels) from Genome in a Bottle (v3.3.2) into the human reference genome sequence (hg38; FASTA format). To minimise computational resources for resulting benchmark experiments, we restricted this to chr22. The commands used were as follows:

```
bcftools consensus --haplotype 1 -f chr22.fa giab_na12878.vcf > hap1.fa

bcftools consensus --haplotype 2 -f chr22.fa giab_na12878.vcf > hap2.fa

cat hap1.fa hap2.fa >  na12878_chr22.fa
```

These commands generate two separate chr22 reference sequences with variants incorporated from NA12878 haplotype 1 and haplotype 2, respectively (homozygous variants are incorporated into both references). We then used *Squigulator* to generate simulated nanopore signal data from this custom diploid reference. To match the data to the NA12878 experimental dataset, we used the *-x dna-r9-prom* pre-set parameter configuration. We adjusted the read-length mean, read-length standard deviation and sequencing depth so as to approximate the equivalent metrics measured from the experimental dataset. The command used was as follows:

```
squigulator na12878_chr22.fa -o reads.blow5 -n 135000 -r 10800 -x dna-r9-prom -t 8 -K 4096
```

***Details of analysis workflow and evaluation with RTG***

Signal data was basecalled with ONT's Guppy software (using the Buttery-eel wrapper for SLOW5 data access; Buttery-eel v0.0.1 on Guppy v6.0.6). Basecalled reads were aligned to the hg38 reference genome with no alternate contigs using Minimap2 (v2.17). Alignment statistics were derived with *Samtools stats* (v1.9). Reference:read identity scores were retrieved using *Paftools*, which is a companion tool in the *Minimap2* repository:

```
samtools view reads.bam -h chr22 | paftools.js sam2paf -p - | awk '{print $10/$11}'
```

Variant calling was performed separately using Nanopolish (v0.14.0) and Clair3 (v0.1-r11; r941_prom_sup_g5014). Variant evaluation was performed using *rtg vcfeval* against the GIAB NA12878 high confidence truth-set (the same callset that was used during the simulation) with QUAL field as the --vcf-score-field. The commands used for basecalling, alignment, variant calling and evaluation were as follows:

```
buttery-eel -i reads.blow5 -o reads.fastq --guppy_bin ont-guppy-6.0.6/bin --port 5887 --config
dna_r9.4.1_450bps_${MODEL}_prom.cfg  -x cuda:all --chunk_size 1500 --max_queued_reads 1000 #
MODEL is fast or hac or sup

minimap2 -x map-ont -a -t32 --secondary=no hg38noAlt.fa reads.fastq  > reads.sam
$SAMTOOLS sort -@32 reads.sam > reads.bam
$SAMTOOLS index reads.bam

run_clair3.sh --threads=32 --include_all_ctgs --bam_fn=reads.bam --ref_fn=hg38noAlt.fa  --platform=ont
--model_path=r941_prom_sup_g5014/ --output=out/ --sample_name=reads --enable_phasing --
longphase_for_phasing
```

```
nanopolish variants -o output.vcf -w ${1}  -r  reads.fastq -g hg38noAlt.fa -b reads.bam -p 2 -t 4 -q cpg --fix-homopolymers

rtg RTG_MEM=32G vcfeval  -b highconf_PGandRTGphasetransfer.vcf.gz  -c merge_output.vcf.gz  -t hg38noAlt.sdf -o compare_clair --region chr22:1-50818468  -e highconf_nosomaticdel_noCENorHET7.bed  --vcf-score-field QUAL
```

***Details of parameter exploration experiment***

For the parameter exploration experiments presented in **Fig3** and **FigS2**, we repeated the simulation and analysis workflows described above, each time varying the simulation parameters. We independently varied the dwell-time mean (--dwell-mean), dwell-time standard deviation (--dwell-std) and amplitude noise factor (--amp-noise), whilst holding the other parameters at the default value. Example commands are as follows:

```
squigulator na12878_chr22.fa -o reads.blow5 -n 135000 -r 10800 -t 8 -K4096  -x dna-r9-prom  --amp-noise <FACTOR> --dwell-mean <MEAN> --dwell-std <STD>
```

For each simulation, the analysis workflow and evaluation was described exactly as above.

***Details for DeepSimulator comparison***

*DeepSimulator* 1.5 main branch on Github (https://github.com/liyu95/DeepSimulator) has an install.sh script for building a conda environment and setting up various other tools required. This script does not work with conda v4+ and thus modifications were made to successfully install *DeepSimulator*. Similarly, the deepsimulatr.sh script for running the *DeepSimulator* pipeline needed modifications to work with conda v4+. Basecalling was excluded from the pipeline when running benchmarks.
To generate simulated libraries for comparison with *Squigulator*, the following commands were run:

```
## for context-independent mode:
deep_simulator.sh -i na12878_chr22_1.fa -o chr22_1_context_ind -n 67500 -l 10800 -c 16
deep_simulator.sh -i na12878_chr22_2.fa -o chr22_2_context_ind -n 67500 -l 10800 -c 16
## for context-dependent mode:
deep_simulator.sh -i na12878_chr22_1.fa -o na12878_chr22_1_context_dep -n 67500 -l 10800 -M 0
deep_simulator.sh -i na12878_chr22_2.fa -o na12878_chr22_2_context_dep -n 67500 -l 10800 -M 0
```

The modified deep_simulator.sh scripts can be found here: [https://github.com/Psy-Fer/DeepSimulator_benchmark](https://github.com/Psy-Fer/DeepSimulator_benchmark)

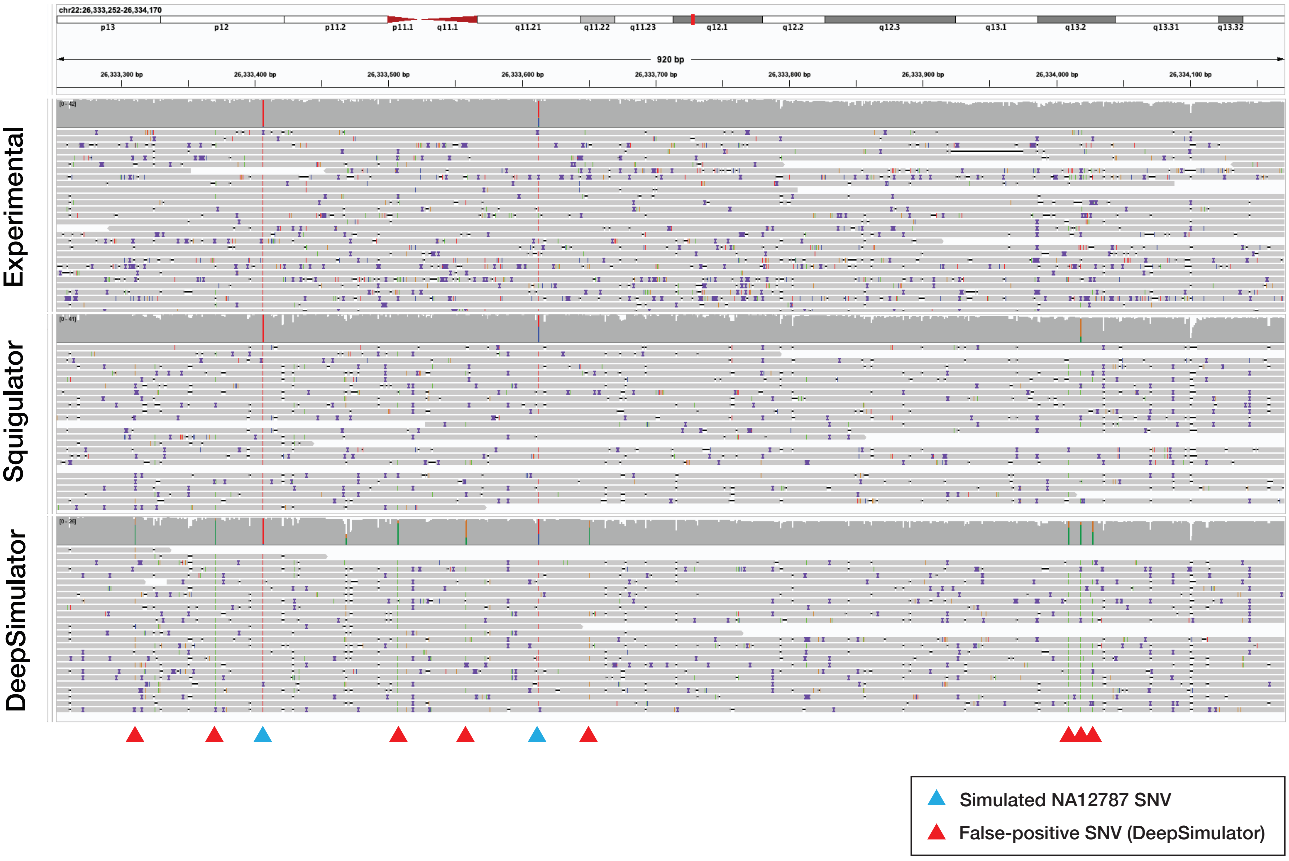


**FigS1. Comparison of Squigulator and DeepSimulator to real experimental ONT data.** Genome browser view shows basecalled reads (Guppy SUP model) aligned to the human reference genome (hg38). The top track shows real experimental data from ONT sequencing of NA12878 genomic DNA (R9.4.1 PromethION flow cells). The middle track shows simulated NA12878 data from Squigulator with -x dna-r9-prom pre-set configuration. The bottom track shows simulated NA12878 data from DeepSimulator running in context-independent mode. Blue triangle markers show the location of NA12878 SNVs that were incorporated into the simulation, and are correctly detected by Clair3. Red triangle markers show the presence of systematic errors in basecalled reads from DeepSimulator, which are erroneously detected as SNVs by Clair3.

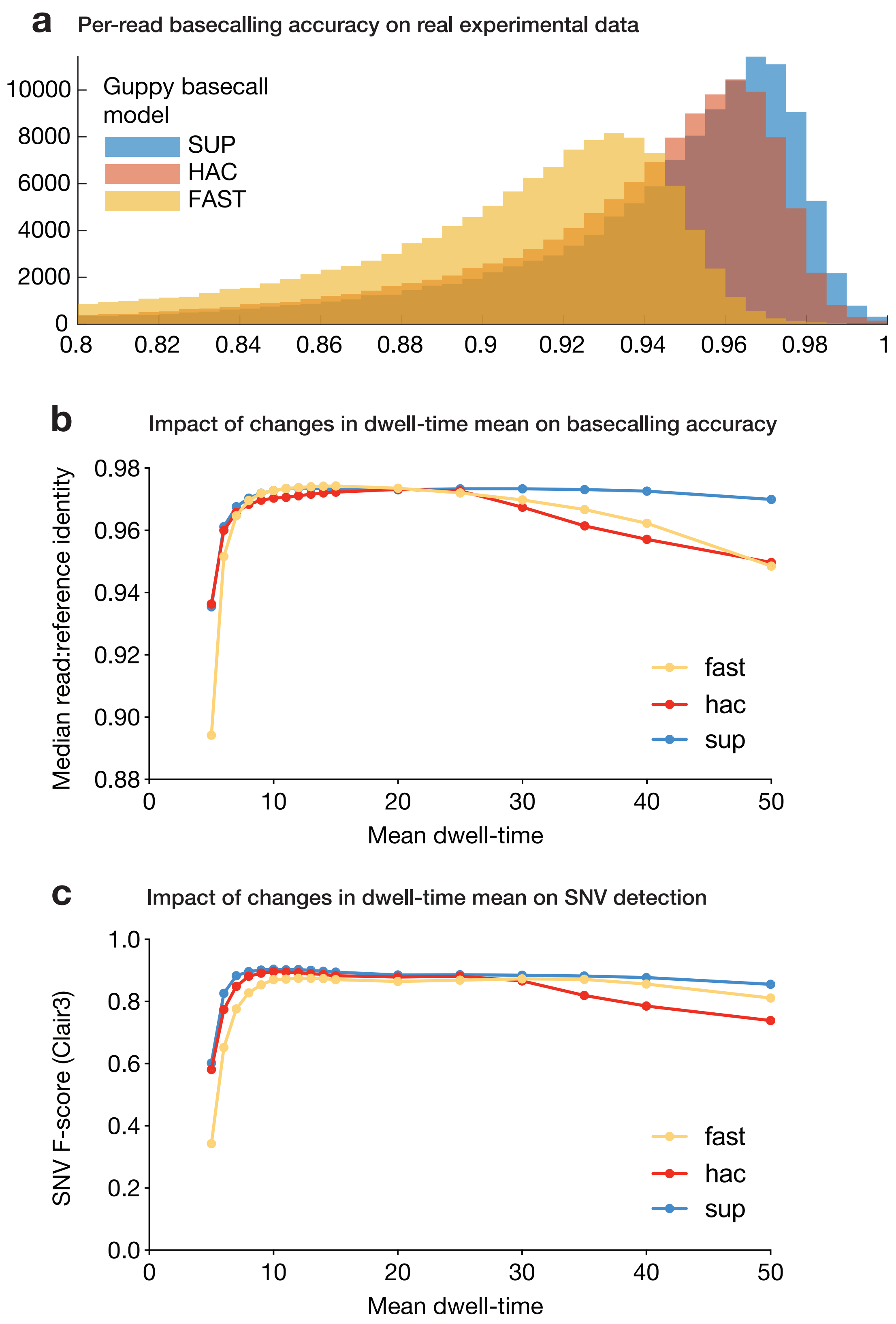


**FigS2. Parameter exploration regarding Guppy basecalling sequencing accuracy.** (**a**) Guppy basecalling accuracy, as measured by read:reference identity score distributions, on real experimental NA12878 data with Guppy's FAST, HAC or SUP models. (**b**) Guppy basecalling accuracy, as measured by read:reference identity score medians, for repeated experiments in which the mean dwell time (--dwell-mean) is varied, while other parameters are held at default. Experiment was repeated with FAST, HAC and SUP basecalling models. Default value --dwell-mean=9 (for R9.4.1 flow cell). (**c**) Accuracy of SNV detection, as measured by F-score, by Clair3 on the same datasets and basecalling models as above (colours are matched).

**Supplementary Table 1: Comparison of Minimap2 alignment statistics for experimental vs simulated NA12878 datasets.**
Basecalled data was generated using *Guppy* HAC model.

| | **NA12878 experimental** | **NA12878 simulated** |
|---|---|---|
| sequences | 135,083 | 134,999 |
| reads mapped | 134,001 | 134,987 |
| reads unmapped | 1,082 | 12 |
| reads MQ0 | 661 | 153 |
| total length | 1,458,924,348 | 1,430,013,633 |
| bases mapped (cigar) | 1,491,898,469 | 1,430,001,381 |
| mismatches | 154,930,196 | 76,290,809 |
| error rate | 1.04E-01 | 5.34E-02 |
| average length | 10800 | 10592 |
| maximum length | 187345 | 87341 |
| average quality | 20.1 | 18 |
| insertions (1-base) | 11,508,645 | 10,218,115 |
| deletions (1-base) | 16,023,165 | 16,965,569 |

**Supplementary Table 2: Comparison of Clair3 and Nanopolish SNV detection statistics for experimental vs simulated NA12878 datasets**
Basecalled data was generated using *Guppy* SUP model.

| Data | Variant caller | Score threshold | True positives baseline | False positives | True positives call | False negatives | precision | sensitivity | f_measure |
|---|---|---|---|---|---|---|---|---|---|
| **Experimental** | **Clair3** | None | 34302 | 118 | 34304 | 160 | 0.9966 | 0.9954 | 0.996 |
| | | 2.15 | 34302 | 118 | 34304 | 160 | 0.9966 | 0.9954 | 0.996 |
| | **Nanopolish** | None | 32743 | 1532 | 32735 | 1719 | 0.9553 | 0.9501 | 0.9527 |
| | | 20.9 | 32713 | 1479 | 32705 | 1749 | 0.9567 | 0.9492 | 0.953 |
| **Simulated** | **Clair3** | None | 33881 | 403 | 33883 | 581 | 0.9882 | 0.9831 | 0.9857 |
| | | 6.48 | 33804 | 255 | 33807 | 658 | 0.9925 | 0.9809 | 0.9867 |
| | **Nanopolish** | None | 33418 | 317 | 33409 | 1044 | 0.9906 | 0.9697 | 0.98 |
| | | 21.7 | 33418 | 316 | 33409 | 1044 | 0.9906 | 0.9697 | 0.9801 |

**Supplementary Table 3: Comparison of Squigulator and DeepSimulator run-time and memory usage.**

Run times and peak RAM usage were measured during simulations of NA12878 data from chr22 at ~30X using 16 CPUs.

| | Squigulator | Deep Simulator (context independent) | Deep Simulator (context dependent) |
|---|---|---|---|
| **Execution time** | 156.194 seconds | 3939.58 seconds | 130.8 hours |
| **Peak RAM usage** | 0.477 GB | 2.117 GB | 49.39 GB |

#### SUPPLEMENTARY FIGURE LEGENDS

**FigS1. Comparison of Squigulator and DeepSimulator to real experimental ONT data**. Genome browser view shows basecalled reads (*Guppy* SUP model) aligned to the human reference genome (hg38). The top track shows real experimental data from ONT sequencing of NA12878 genomic DNA (R9.4.1 PromethION flow cells). The middle track shows simulated NA12878 data from *Squigulator* with *-x dna-r9-prom* pre-set configuration. The bottom track shows simulated NA12878 data from *DeepSimulator* running in context-independent mode. Blue triangle markers show the location of NA12878 SNVs that were incorporated into the simulation, and are correctly detected by *Clair3*. Red triangle markers show the presence of systematic errors in basecalled reads from *DeepSimulator*, which are erroneously detected as SNVs by *Clair3*.

**FigS2. Parameter exploration regarding *Guppy* basecalling sequencing accuracy.** (**a**) Guppy basecalling accuracy, as measured by read:reference identity score distributions, on real experimental NA12878 data with *Guppy's* FAST, HAC or SUP models. (**b**) Guppy basecalling accuracy, as measured by read:reference identity score medians, for repeated experiments in which the mean dwell time (--dwell-mean) is varied, while other parameters are held at default. Experiment was repeated with FAST, HAC and SUP basecalling models. Default value --dwell-mean=9 (for R9.4.1 flow cell). (**c**) Accuracy of SNV detection, as measured by F-score, by *Clair3* on the same datasets and basecalling models as above (colours are matched).
